# Rescue of Impaired Blood-Brain Barrier in Tuberous Sclerosis Complex Patient Derived Neurovascular Unit

**DOI:** 10.1101/2023.12.15.571738

**Authors:** Jacquelyn A. Brown, Shannon L. Faley, Monika Judge, Patricia Ward, Rebecca A. Ihrie, Robert Carson, Laura Armstrong, Mustafa Sahin, John P. Wikswo, Kevin C. Ess, M. Diana Neely

**Author notes:** Co-Senior and Corresponding authors.

## Abstract

Tuberous sclerosis complex (TSC) is a multi-system genetic disease that causes benign tumors in the brain and other vital organs. The most debilitating symptoms result from involvement of the central nervous system and lead to a multitude of severe symptoms including seizures, intellectual disability, autism, and behavioral problems. TSC is caused by heterozygous mutations of either the *TSC1* or *TSC2* gene. Dysregulation of mTOR kinase with its multifaceted downstream signaling alterations is central to disease pathogenesis. Although the neurological sequelae of the disease are well established, little is known about how these mutations might affect cellular components and the function of the blood-brain barrier (BBB). We generated disease-specific cell models of the BBB by leveraging human induced pluripotent stem cell and microfluidic cell culture technologies. Using these microphysiological systems, we demonstrate that the BBB generated from *TSC2* heterozygous mutant cells shows increased permeability which can be rescued by wild type astrocytes and with treatment with rapamycin, an mTOR kinase inhibitor. Our results further demonstrate the utility of microphysiological systems to study human neurological disorders and advance our knowledge of the cell lineages contributing to TSC pathogenesis.

## Background

Neurogenetic disorders often present in young children due to their impact on the developing brain. The neurological manifestations of such disorders are typically severe and include epilepsy, intellectual disabilities, and autism (Moffat, Ka et al. 2015). Tuberous Sclerosis Complex (TSC) has been a prototypical model for neurogenetic disease for many years (Curatolo, Bombardieri and Jozwiak 2008). The study of TSC provides several advantages, including well-described clinical manifestations and a known genetic etiology from loss of *TSC1* or *TSC2* gene function with defined downstream signaling pathways (Ess 2006, Rosset, Netto and Ashton-Prolla 2017, Rosset, Vairo et al. 2017). In addition, affected downstream signaling pathways have been identified, with an ostensibly central role for dysregulation of mTOR kinase (Baybis, Yu et al. 2004, Kaper, Dornhoefer and Giaccia 2006, Lim, Gopalappa et al. 2017). Hamartin and tuberin, the *TSC1* and *TSC2* protein products respectively, normally inhibit mTOR signaling through an indirect pathway. *TSC1* and *TSC2*-mutant cells thus have constitutively increased activity of mTOR kinase, which likely underlies the abnormal proliferation and differentiation of cells suspected to occur in multiple organs of patients with TSC (Ihrie and Henske 2022). This understanding has led to the development of mTOR inhibitors (“rapalogs”) that are FDA-approved and increasingly used for the treatment of several aspects of TSC (Krueger, Care et al. 2010, McCormack, Inoue et al. 2011, Bissler, Kingswood et al. 2013, Krueger, Capal et al. 2018).

While many animal models (mouse, rat, zebrafish, *Drosophila*) have been developed over the past 20 years (Uhlmann, Wong et al. 2002, Wong, Ess et al. 2003, Wenzel, Patel et al. 2004, Meikle, Talos et al. 2007, Kim, Speirs et al. 2011, Carson, Van Nielen et al. 2012, Tsai, Hull et al. 2012, Carson, Fu et al. 2013, Carson, Kelm et al. 2015), many key aspects of TSC pathogenesis remain poorly understood. These include species-specific impact of mTOR signaling as well as the requirements for heterozygous versus homozygous mutations of the *TSC1*/*TSC2* genes. The advance of human induced pluripotent stem cell (iPSC) technology (Dolmetsch and Geschwind 2011, Higurashi, Uchida et al. 2013, Nityanandam and Baldwin 2015, Snow, Westlake et al. 2020) combined with tissue chip technology (Brown, Pensabene et al. 2015, Brown, Codreanu et al. 2016, Vernetti, Gough et al. 2017, Brown, Faley et al. 2020) has allowed the use of human tissue-based models to address these fundamental questions.

Although the neurological sequelae of TSC are well established, little is known about how *TSC1* or *TSC2* mutations affect different cellular and functional components of the brain. While expression of the astrocytic protein aquaporin-4, a component of the blood-brain barrier (BBB) is increased in epileptic cortex from patients with TSC and TSC mouse models (Short, Kozek et al. 2019), BBB function has not been well studied in TSC. We hypothesized that an abnormal BBB contributes to TSC pathogenesis. The study of the BBB in animal models of TSC although highly valuable also presents some challenges. Thus, species-specific differences in human versus mouse brain structure and cellular function have recently become more apparent. In addition, complicated genetic manipulations are needed to study the impact of *Tsc2* mutations in specific cell types. Finally, the frequent occurrence of epilepsy or brain tumor phenotypes in animal models also represent potential confounding factors, as these symptoms could have secondary effects on the BBB. Therefore, to test our hypothesis and examine primary effects of *TSC2* mutation on the BBB, we used an *in vitro* neurovascular unit (NVU) (**Figure 1**) to create TSC patient-specific brain tissue models that were generated by leveraging iPSC and microfluidic cell culture technologies (Brown, Pensabene et al. 2015, Brown, Codreanu et al. 2016, Brown, Faley et al. 2020). Using these microphysiological systems, we demonstrate that the *TSC2* patient derived BBB shows increased permeability which can be rescued with rapamycin, an mTOR kinase inhibitor. Astrocytes appear to play a major role in this TSC BBB phenotype as replacement of *TSC2*-mutant astrocytes with wild type astrocytes also rescued the TSC2 patient derived BBB defect.

**Figure 1.**
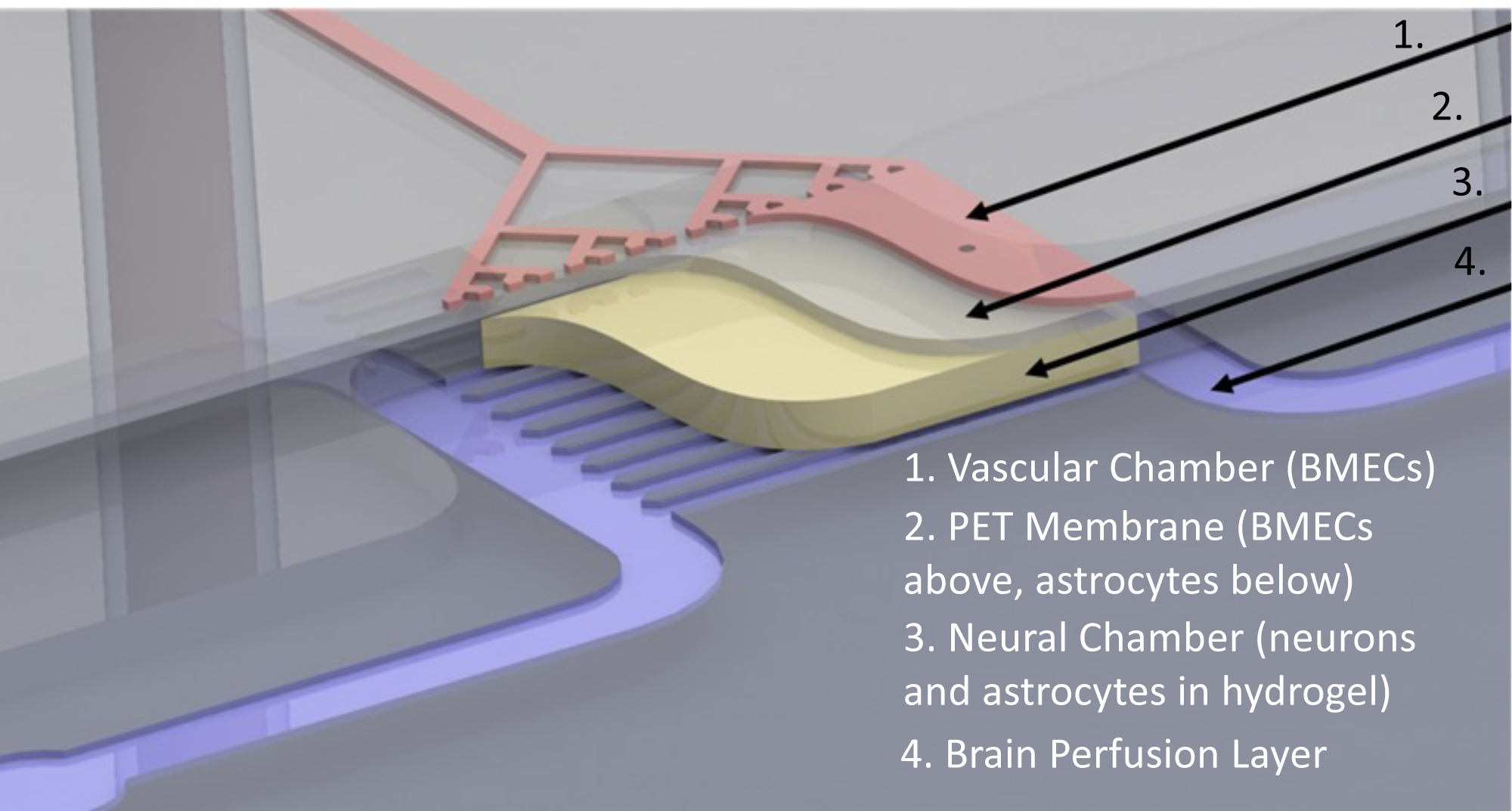
Schematic overview of NVU. The vascular chamber **(1, Pink)** into which BMECs are loaded and the media for the vascular chamber is perfused. Porous PET membrane **(2, Grey)** with 3 µm pores supports a layer of BMECs on one side and a layer of astrocytes on the opposite “brain” neural cell chamber side. Neural chamber **(4, Yellow)** contains neurons and additional astrocytes within a hydrogel. Perfusion channels through the neural cell chamber indicated as **Purple**.

Our findings substantiate the use of microphysiological systems to study neurogenetic disorders. We interpret our results within what is known about the BBB in TSC and discuss how our findings expand this knowledge and support ongoing translational research related to TSC pathogenesis and treatment.

## Methods

### Derivation, validation, and differentiation of iPSCs

Three of the human induced pluripotent stem cell (iPSC) lines (control CC3, TSC patient TSP8-15, and TSP23-9) were derived at Vanderbilt University Medical Center and validated according to established protocols (Armstrong, Westlake et al. 2017, Neely, Davison et al. 2017). In brief, primary dermal fibroblasts were established from skin biopsies obtained after patients’ consent/assent under the Vanderbilt University Medical Center IRB protocol #080369. Fibroblasts were reprogrammed by electroporation with either CXLE plasmid vectors (Howden, Thomson and Little 2018) (lines CC3 and TSP8-15) using the Neon Transfection System (Life Technologies, Carlsbad, CA, USA) or CytoTune iPS 2.0 Sendai Reprogramming Kit (ThermoFisher) (line TSP23-9). Transfected fibroblasts were then plated at 5 x 10^4^ cells/well into Matrigel-coated 6-well plates. Two days later, cells were transferred into TeSR-E7 medium and maintained until the emergence of iPSC colonies (about 4 weeks) which were then manually isolated and propagated in mTeSR medium (StemCell Technologies). Absence of plasmid integration and clearance of Sendai virus were confirmed, and normal karyotype verified using at least 20 metaphase spreads (Genetics Associates, Nashville, TN, USA) (**Supplementary** Figure 1A). Pluripotency markers were present as previously reported and pluripotency was further validated by Pluritest and/or the ability of the iPSC lines to differentiate into all three germline lineages using the hPSC TaqMan Scorecard (ThermoFisher A15870), and into neural lineages as we have previously described (Neely, Tidball et al. 2011, Brown, Codreanu et al. 2016, Tidball, Neely et al. 2016, Neely, Davison et al. 2017, Neely, Xie et al. 2021, Prince, Neely et al. 2021) (Neal, Marinelli et al. 2019). Isogenic iPSC lines TSP77+/+ and TSP77+/− were generated at Boston’s Children Hospital and previously validated as described (Sundberg, Tochitsky et al. 2018).

### Cortical glutamatergic neuron differentiation

iPSCs were replated at a density of 2 x 10^4^ cells/cm^2^ in mTeSR medium containing 10 µM Rho kinase inhibitor (Y-27632, Tocris #5849), which was removed after 24 hours. Once the cultures reached 100% confluency (day 0), neuralization was induced via an eleven-day dual-SMAD inhibition protocol using 0.4 µm LDN (Tocris # 6053) and 10 µM SB 431542 (Tocris #1614) as previously described (Chambers, Fasano et al. 2009, Neely, Litt et al. 2012). Starting on day 11, the neuronal cultures underwent further differentiation in cortical differentiation medium as previously reported (Neely, Xie et al. 2021, Prince, Neely et al. 2021). Around day 20-25, differentiating cells were passaged for the first time by incubating them with Accutase (StemCell Technologies, #01-0006) for 18-30 minutes and reseeding them into Matrigel (BD Bioscience #354277)-coated 6-well plates at 1 x 10^5^ cells/cm^2^ in cortical differentiation medium containing 10 µM Rock-inhibitor (Tocris, Minneapolis, MN, USA #1254), which was removed after 24 hours. Cultures were maintained in cortical differentiation medium and replated at 3 x 10^5^ cells/cm^2^ monthly. Neurons were harvested for seeding into the neurovascular units (NVUs) between days 80-120 of differentiation from iPSC stage. The cortical cultures derived from all iPSC lines contained abundant neurons with dense and complex neurite projections (**Supplementary** Figures 2A and B).

### Cortical astrocyte differentiation

We used a “spontaneous emergence approach” (Chandrasekaran, Avci et al. 2016) to make astrocyte cultures, as we have previously described (Miller, Schaffer et al. 2021). By around day 80 of cortical glutamatergic neuron differentiation, we begin to see astrocytes emerging alongside cortical glutamatergic neurons. By day 120-160 of differentiation, we passaged cells monthly at low density (0.5 x 10^5^ cells/cm^2^), which leads to a continuous loss of neurons from the cultures and results in relatively pure astrocyte cultures after 3-4 passages. At that point astrocytes are transferred into astrocyte medium (ScienCell #1801) and replated at 0.4 x 10^5^ cells/cm^2^ monthly until integration into the NVU. The large majority of cells in these astrocyte cultures express the astrocyte marker GFAP and/or S100B (**Supplementary** Figures 3).

### BMEC-like differentiation

Brain microvascular endothelial-like cells (BMECs) were differentiated from iPSCs according to previously reported protocols, with minor modifications (Bosworth, Faley et al. 2017, Faley, Neal et al. 2019). Briefly, iPSCs were seeded at a concentration of 150,000 cells per well of a Matrigel-coated 6-well plate in E8 medium (ThermoFisher) containing Rock-inhibitor (10 µM; Y-27632). The following day, E8 medium was replaced with E6 medium (ThermoFisher Scientific) and changed daily for 4 days. On day 4, the medium was changed to Neurobasal (Gibco) medium supplemented with B27 (Gibco), 0.5 mM Glutamax (ThermoFisher Scientific), 20 ng/ml basic fibroblast growth factor (bFGF; PeproTech, Rocky Hill, NJ, USA), and 10 μM all-trans retinoic acid (RA; ThermoFisher Scientific) for 48 hours. BMECs were subcultured on day 6 in the same medium containing Rock-inhibitor (10 µM Y-27632). For traditional experiments, BMECs were subcultured either onto Polyester PET Transwells of 3 µm pore size or into 12-well tissue culture plates (Corning), both of which had been coated with a mixture of collagen IV (Sigma-Aldrich, 400 ng/ml) and fibronectin (Sigma-Aldrich, 100 ng/ml) in PBS for 24 hours at 37°C.

### Transendothelial electrical resistance (TEER) Measurements

TEER was measured using an EVOM voltohmmeter with STX2 electrodes (World Precision Instruments) using standard protocols we have previously employed (Brown, Pensabene et al. 2015, Bosworth, Faley et al. 2017, Faley, Neal et al. 2019, Brown, Faley et al. 2020). Raw TEER values were adjusted through subtracting TEER values measured across an empty filter and then multiplied by filter surface area to yield TEER (Ωxcm^2^).

### Transwell culture

Transwell cultures were seeded and cultured in parallel with the NVU cultures to allow for a direct comparison of cell phenotypes in these two different models. Thus, Transwell plates were coated with the same ECM components used in the vascular side of the NVU and seeded with the same BMECs suspension in the same medium. The BMECs were seeded onto the membrane of the apical side of the Transwell insert. Sample collection from the NVU and Transwell cultures was also performed in parallel.

### NVU cell load and culture

NVU devices were coated with collagen IV and fibronectin as previously described (Brown, Pensabene et al. 2015). BMECs (control or *TSC2*-mutant) were seeded into the vascular chamber at a density of (8–10)L×L10^6^ cells/ml (day 0) (Brown, Pensabene et al. 2015, Brown, Codreanu et al. 2016). The device was oriented (neural cell “brain” chamber bottom, vascular chamber top) to allow BMEC attachment to the membrane overnight. On the following day, the astrocytes were loaded into the neural cell chamber at (2-5)L×L10^6^ cells/ml and the devices were inverted (neural cell chamber top, vascular chamber bottom) to allow for attachment of the astrocytes on the membrane side opposite to the seeded BMEC. Two hours after astrocyte loading, neurons (8–10)L×L10^6^ cells/ml suspended in a hydrogel (Mebiol Gel; Cosmo Bio Co. Ltd, #MBG-PMW20) were seeded into the neural cell chamber as previously described (Brown, Pensabene et al. 2015, Brown, Codreanu et al. 2016). Once the hydrogel had set, typically after 1 hour, media flow was started. The vascular chamber containing BMECs was perfused with Lonza EGM-2 medium and the neural chamber containing neurons and astrocytes was perfused with Neurobasal medium supplemented with B27 and Glutamax. Devices were maintained for a minimum of 24–48 hours before experiments were conducted.

### NVU BBB permeability assay

#### Passive permeability measurements in NVUs

To measure BBB permeability in the NVUs we added 3kD (unless indicated otherwise) Alexa Fluor 680 dextran (excitation 680 and emission 706 nm; ThermoFisher) at a final concentration of 1 µM to the vascular compartment medium and perfused the NVU with this dextran solution for the entire duration of the culture (Brown, Faley et al. 2020). For passive permeability measurements, effluent was collected from the neural cell compartment over a fixed amount of time and the fluorescence intensity of the eluate quantified using a plate reader (Tecan M1000). Using the determined dye concentration in the brain compartment, we calculated the applied permeability coefficient, *P_app,_* using the standard equation:

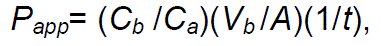

where *V_b_* is the neural cell chamber volume in cm^3^, *A* is the vascular chamber growth area in cm^2^, *C_a_* is the dextran concentration perfused into in the vascular compartment (µM), *C_b_* is the dextran concentration (µM) measured in the neural cell compartment eluate, and *t* is the assay time in seconds (Frost, Jiang et al. 2019).

The effective permeability of the BMEC monolayer, *P_cells_*, was calculated from the measured applied permeability P_app_ by correcting for the permeability of a device without cells (but containing hydrogel and a membrane coated with collagen/fibronectin), *P_membrane_*, according to the equation:

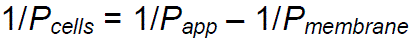

### Rapamycin treatment

Rapamycin (Thermo-Fisher) was reconstituted as a 10 µM stock solution in DMSO. 10 nM rapamycin in culture medium was then either perfused through the vascular chamber of the NVUs or added to the apical chamber of the Transwell cultures for 24 hours, after which BBB permeability measurements were carried out as described above.

### Metabolomics

Neural cell and vascular chamber samples were stored at –80°C until analyzed via Liquid Chromatography-High Resolution Mass Spectrometry (LC-HRMS)-based metabolomics in the Vanderbilt Center for Innovative Technology (CIT) using previously described methods (Brown, Codreanu et al. 2016, Rogers, Sobolik et al. 2018, Mancini, Schrimpe-Rutledge et al. 2021) Briefly, effluent samples collected from both the vascular and brain chambers of the NVU were normalized by volume to 100 uL as previously reported (Brown, Codreanu et al. 2016). Metabolites were extracted with methanol/water 80:20. Heavy labeled phenylalanine-D8 and biotin-D2 were added to individual samples prior to protein precipitation. Following overnight incubation at –80°C, precipitated proteins were pelleted by centrifugation at 10,000 rpm for 10 min and metabolite extracts were dried down *in vacuo* and stored at –80°C.

Individual extracts were reconstituted in 50 µl of acetonitrile/water (3:97, v/v) with 0.1% formic acid containing heavy-labeled carnitine-D9, tryptophan-D3, valine-D8, and inosine-4N15, and centrifuged for 5 min at 10,000 rpm to remove insoluble material. A pooled quality control sample (QC) was prepared by pooling equal volumes of individual samples. The pooled QC sample was used for column conditioning (8 injections prior to sample analysis), retention time alignment and to assess mass spectrometry instrument reproducibility throughout the sample set.

Global, untargeted mass spectrometry analyses were performed on a high-resolution Q-Exactive HF hybrid quadrupole-Orbitrap mass spectrometer (Thermo Fisher Scientific, Bremen, Germany) equipped with a Vanquish UHPLC binary system (Thermo Fisher Scientific, Bremen, Germany). Extracts (5μL injection volume) were separated on a Hypersil Gold, 1.9 μm, 2.1 mm × 100 mm column (Thermo Fisher) held at 40 °C. LC was performed at 250 μL/min using solvent A (0.1% FA in water) and solvent B (0.1% FA in acetonitrile/water 80:20) with a gradient length of 30 min as previously described (Eberly, Beebout et al. 2020, Popay, Wang et al. 2021). Full MS analyses were acquired over the mass-to-charge ratio (*m/z*) range of 70-1,050 in positive ion mode. Full mass scan was acquired at 120,000 resolution with a scan rate of 3.5 Hz, automatic gain control (AGC) target of 1×10^6^, and maximum ion injection time of 100 ms, and MS/MS spectra were collected at 15,000 resolution, AGC target of 2×10^5^ ions, with a maximum ion injection time of 100 ms.

### Metabolomics Data Processing and Pathway Analysis

Mass spectrometry raw data was imported, processed, normalized and reviewed using Progenesis QI v.3.0 (Non-linear Dynamics, Newcastle, UK). All MS and MS/MS sample runs were aligned against a pooled QC reference run. Unique ions (retention time and m/z pairs) were de-adducted and de-isotoped to generate unique “features” (retention time and m/z pairs). Data were normalized to all features and significance was assessed using p-values generated using ANOVA (analysis of variance) from normalized compound abundance data. Tentative and putative annotations were determined by using accurate mass measurements (<5 ppm error), isotope distribution similarity, and fragmentation spectrum matching (when applicable) by searching the Human Metabolome Database (Wishart, Jewison et al. 2013), METLIN (Smith, O’Maille et al. 2005), and the CIT’s in-house library. Annotations (Confidence Level 1-3 (Schrimpe-Rutledge, Codreanu et al. 2016) were determined for all compounds with a match to any of the searched libraries or databases. Metaboanalyst 5.0 (www.metaboanalyst.ca/) was used to perform pathway and metabolite enrichment analyses from annotated compounds with statistical significance (p-value ≤ 0.05) (Chong, Wishart and Xia 2019).

### RNA Seq

Total RNA was extracted from cells and purified with Rneasy mini kit (Qiagen) and quality was validated with an Agilent kit. RNA sequencing reads were adapter-trimmed and quality-filtered using Trimgalore v0.6.7 (Krueger et al., 2023). An alignment reference was generated from the GRCh38 human genome and GENCODE comprehensive gene annotations (Release 26), to which trimmed reads were aligned and counted using Spliced Transcripts Alignment to a Reference (STAR) v2.7.9a (Dobin et al., 2013) with quantMode GeneCounts parameter. Approximately 60 million uniquely mapped reads were acquired per sample. DESeq2 package v1.36.0 (Love et al. 2014) was used to perform sample-level quality control, low count filtering, normalization, and downstream differential gene expression analysis. Genomic features counted fewer than five times across at least three samples were removed. False discovery rate adjusted for multiple hypothesis testing with Benjamini-Hochberg (BH) procedure *p* value < 0.05 and log2 fold change >1 was used to define differentially expressed genes. Pairwise comparison with *TSC2* heterozygous mutation (TSP8-15) versus control (CC3) was performed. Three replicates per condition were included for the differential gene expression analysis. Gene set enrichment analysis (GSEA) was performed using the R package Clusterprofiler (Yu, Wang et al. 2012) with gene sets from the Human MSigDB database v2022.1.Hs (Subramanian, Tamayo et al. 2005).

### Statistics

Statistical significance was determined using a *t* test when changes were compared between two groups (GraphPad Prism 6). *P*< 0.05 was considered statistically significant unless otherwise stated. Values were expressed as means ± SEM. For comparisons of more than 2 groups, we used either one-way ANOVA (for normal distributions) with Bonferroni post hoc test or Kruska Wallis with Dunn’s post hoc test where a normal distribution cannot be confirmed. (p < 0.01 is indicated with ** in the figures throughout the manuscript).

### Data Availability

The RNA-Seq data has been annotated and deposited to NCBI GEO with accession ID GSE235862.

Metabolomics data is in the process of being submitted to the NIH Common Fund’s National Metabolomics Data Repository (NMDR) website, the Metabolomics Workbench, https://www.metabolomicsworkbench.org. This work is supported by Metabolomics Workbench/National Metabolomics Data Repository (NMDR) (grant# U2C-DK119886), Common Fund Data Ecosystem (CFDE) (grant# 3OT2OD030544) and Metabolomics Consortium Coordinating Center (M3C) (grant# 1U2C-DK119889).

## Results

We initially used conventional monolayers of human iPSC-derived brain microvascular endothelial cells (BMECs) grown in a Transwell dish to evaluate barrier permeability of BMECs differentiated from control (CC3) and *TSC2* heterozygous mutant (TSP8-15) iPSC lines. We delivered fluorescently labelled dextran of different sizes (300 or 3,000 Da) to the apical side of the barriers and quantified subsequent concentration in the basal side. The relative permeability of TSC mutant BMEC barriers showed a trend towards higher permeability than barriers from control BMECs for the 300 Da dextran, but the difference did not reach statistical significance (**Figure 2A**). As expected, pericytes, which do not form tight junctions (Bauer, Krizbai et al. 2014), cannot form a barrier other than the passive resistance created by the presence of cells in a monolayer on the filter membrane. Thus, the permeability for pericytes is greater than that of BMEC cultures (**Figure 2A**). We also looked at transendothelial electrical resistance (TEER) of BMECs cultured on membranes of varying pore sizes to evaluate the integrity of the endothelial barrier. A two-way ANOVA analysis showed that TEER was significantly affected by genotype, but not by membrane pore size (**Figure 2B**).

**Figure 2.**
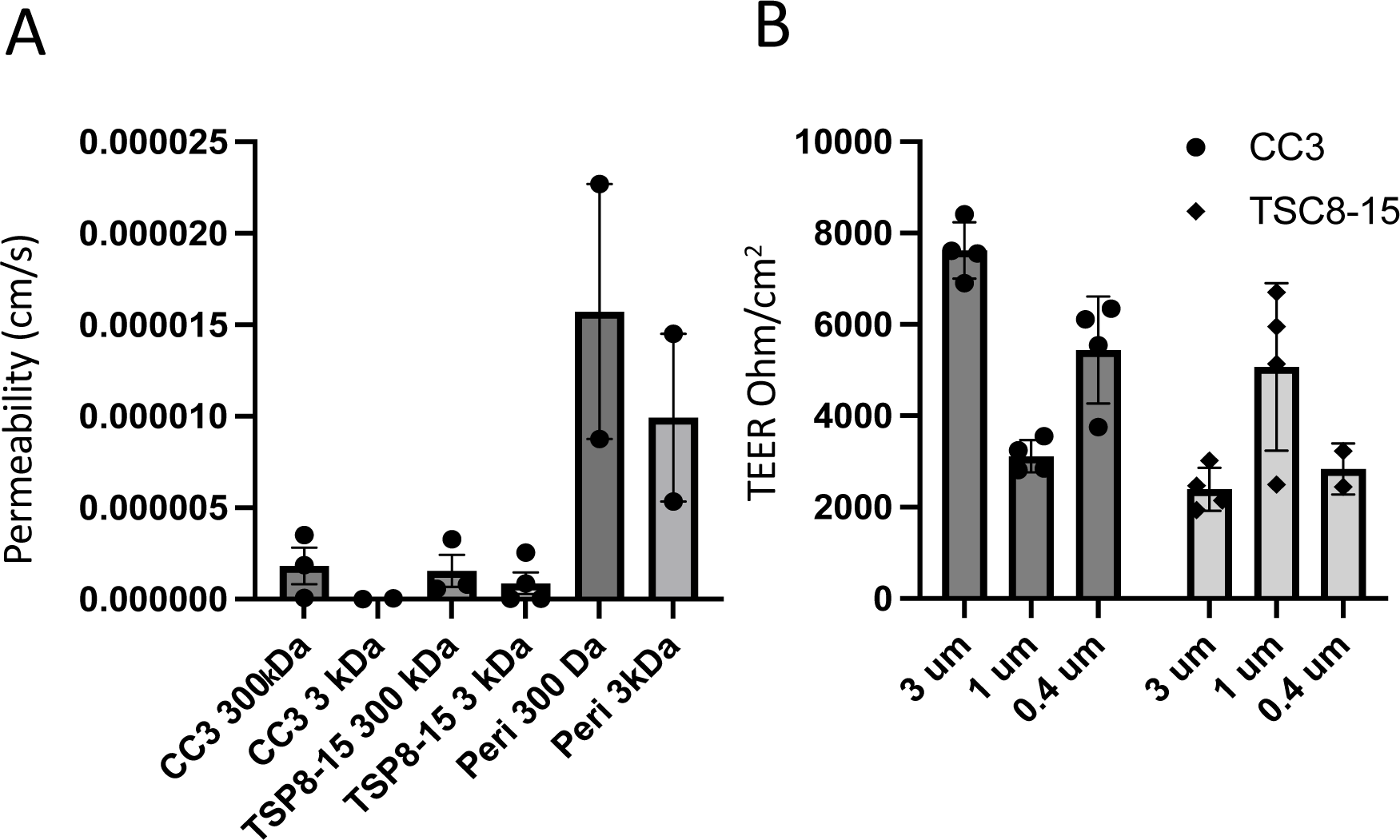
Comparison of barrier function of control and *TSC2*-mutant BMEC monocultures in Transwell plates. A) TSC patient-derived (TSP8-15) BMEC transwell cultures showed a statistically non-significant trend towards larger permeability for the 3000 DA, but not 300 Da dextrans when compared to control (CC3) BMEC cultures. Pericytes (Peri) formed a minimal barrier (N = 2-4). **B)** Transendothelial electrical resistance (TEER) of CC3 and TSP8-15 BMECs cultured on membranes of varying pore sizes were measured. A two-way ANOVA analysis of the data indicated a genotype-dependent F (1,16) = 18.87 (P < 0.0005), but membrane pore-size independent F(2,16) = 1.91 (P = 0.1791) effect on TEER (overall P < 0.0001, N= 2-4).

We next compared barrier permeabilities between transwell BMEC monocultures and the more complex multi-cell NVU assembly with “brain” (neural cell) and “vascular” sides (**Figure 1**) using 3 kD dextran. The difference between CC3 and TSP8-15 barrier function in the monoculture transwell model was not statistically significant, though TSP8-15 derived BMECs trended towards higher permeability (**Figure 3A**). In NVUs seeded with BMECs, astrocytes, and neurons we found a statistically significantly higher permeability in TSP8-15 than in control CC3-cell seeded NVUs (**Figure 3B**) (p < 0.01, N=5). The expression of the BMEC marker proteins occludin, claudin, and ZO-1 validated the BMEC-like nature of our CC3 and TSP8-15 cultures (**Figure 4A**). **Figure 4B** shows phase microscopy images of seeded vascular and neural compartments of CC3 and TSP8-15 NVUs.

**Figure 3.**
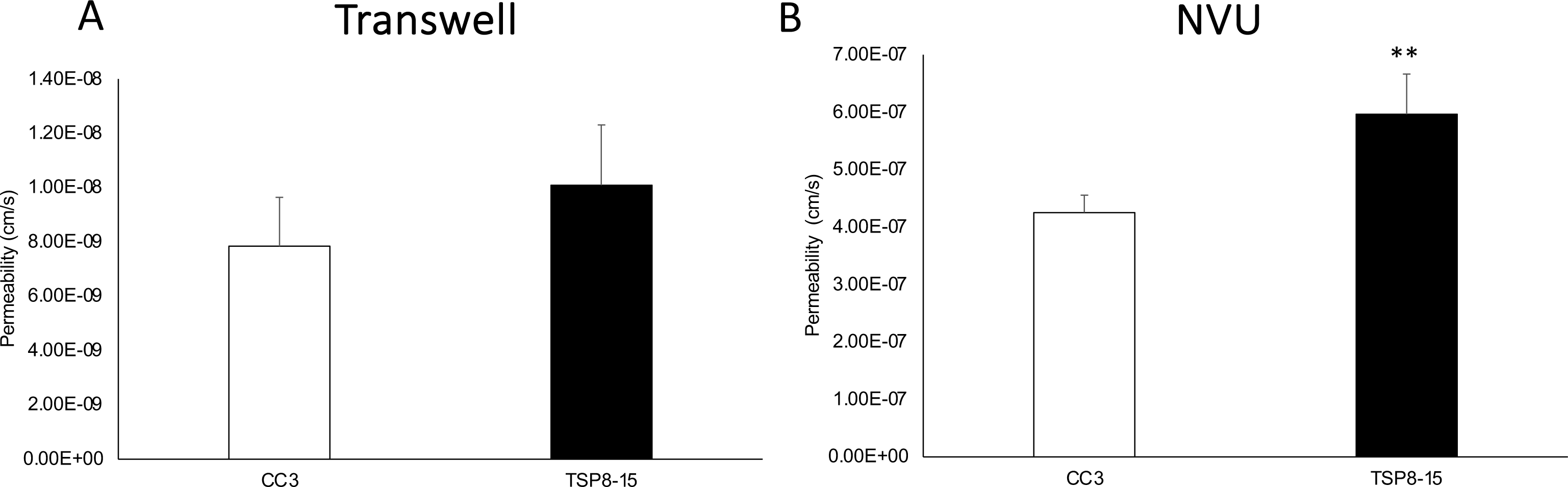
Disruption of BBB in an NVU co-culture system. (A) TSC patient-derived (TSP8-15) BMEC Transwell cultures 8 days in culture show a small, but not statistically significant higher permeability with 3 kD FITC dextran as compared to CC3 (p< 0.06, N=5). **(B)** In the NVU system, after 8 days co-culture with astrocytes and neurons of matching genotype, BBB permeability for the TSP8-15 was significantly increased compared to CC3 (p < 0.01, N=5).

**Figure 4.**
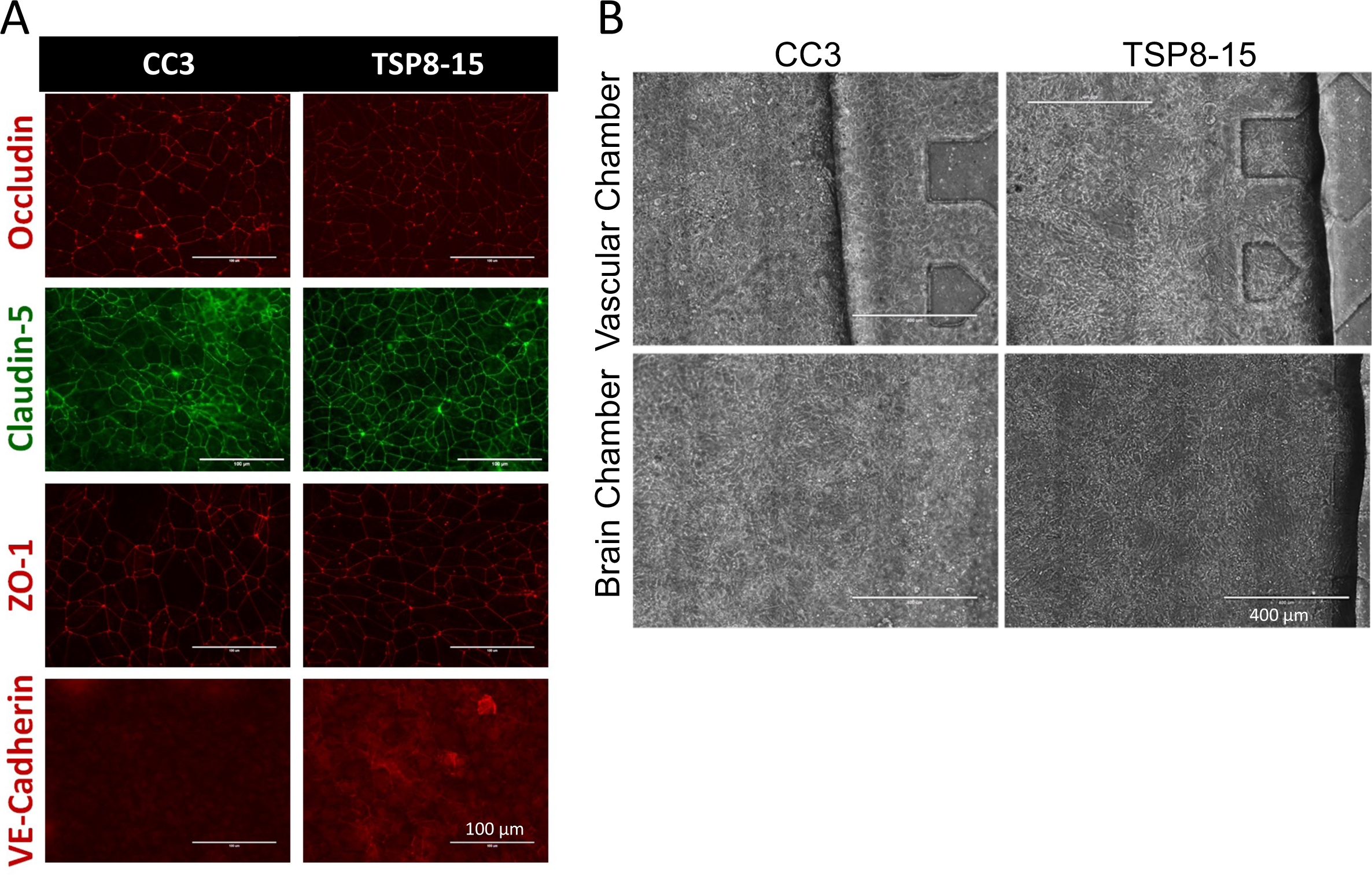
Expression of junctional proteins in BMEC cultures. (A) BMEC monolayers derived from CC3 and TSP8-15 iPSC lines express BMEC markers occludin, claudin-5, and ZO-1 and VE-cadherin. Scale bar for all images = 100 µm. **(B)** Phase images of seeded CC3 and TSP8-15 NVU vascular BMEC-containing compartments (top panels) and the neuron and astrocyte-containing neural cell chambers (bottom panels) are shown. Scale bar in all images = 400 µm.

We extended our BBB permeability analyses to a set of isogenic iPSCs (TSP77; either wild type or heterozygous for a *TSC2* mutation). As seen for CC3 control– (*TSC2* wild type) and TSP8-15 *TSC2* heterozygous mutant line BMECs, TSP77+/+ and TSP77+/− derived BMECs express the typical BMEC markers claudin-5, occluding, ZO-1 and Glut-1 (**Figure 5A**). Importantly, we observed a significantly higher permeability in TSP77+/− *TSC2* heterozygous mutant cultures compared to TSP77+/+ isogenic wild type cultures (**Figure 5B**), thus confirming our permeability observations in the CC3 and TSP8-15 NVUs (**Fig. 3B**). A trend towards a higher difference in TSC (TSP8-15) versus control (CC3) BBB permeability was observed as early as day 2 after NVU seeding, and was significant by day 5 and thereafter (**Fig. 6A and Fig. 7A**).

**Figure 5.**
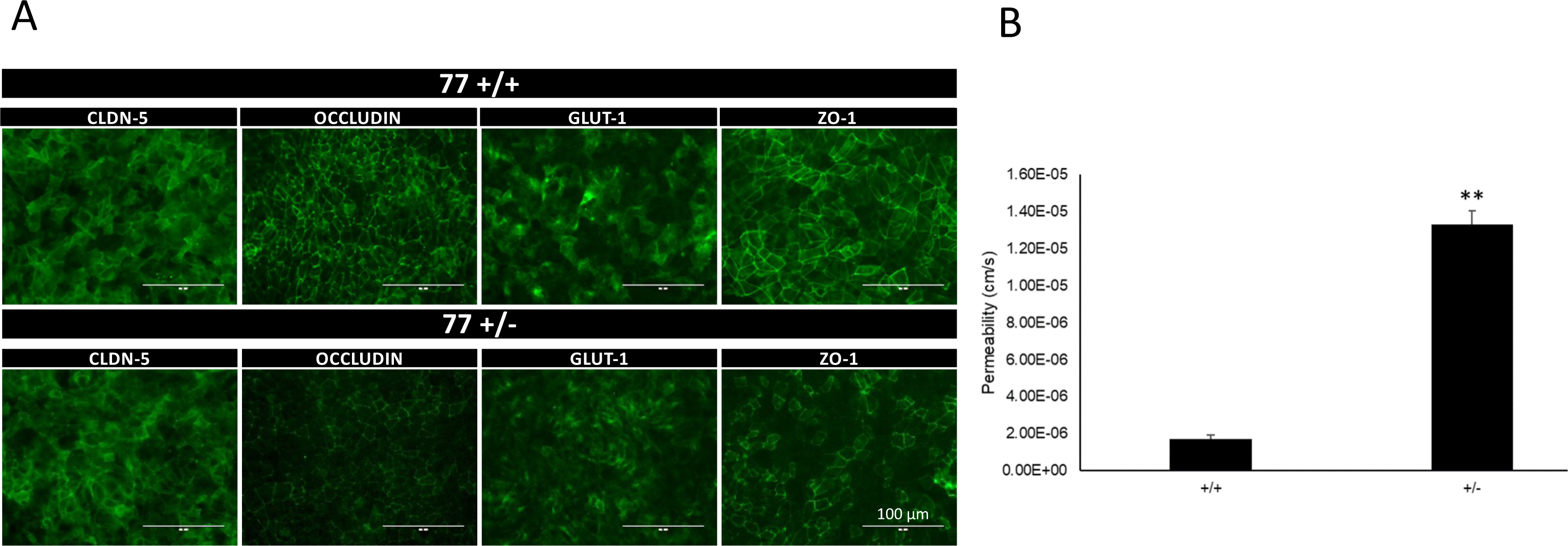
Increased BBB permeability in NVU generated from *TSC2* heterozygous mutant cells compared to isogenic control cells. (A) TSP77+/+ and TSP77+/− derived BMECs express the BMEC marker proteins occluding, claudin-5, ZO-1 and Glut-1. Scale bar for all images 100 µm. **(B)** The BBB permeability in TSP77+/− NVUs is significantly increased compared to isogenic control TSP77 +/+ NVUs (p < 0.01, N = 5).

**Figure 6.**
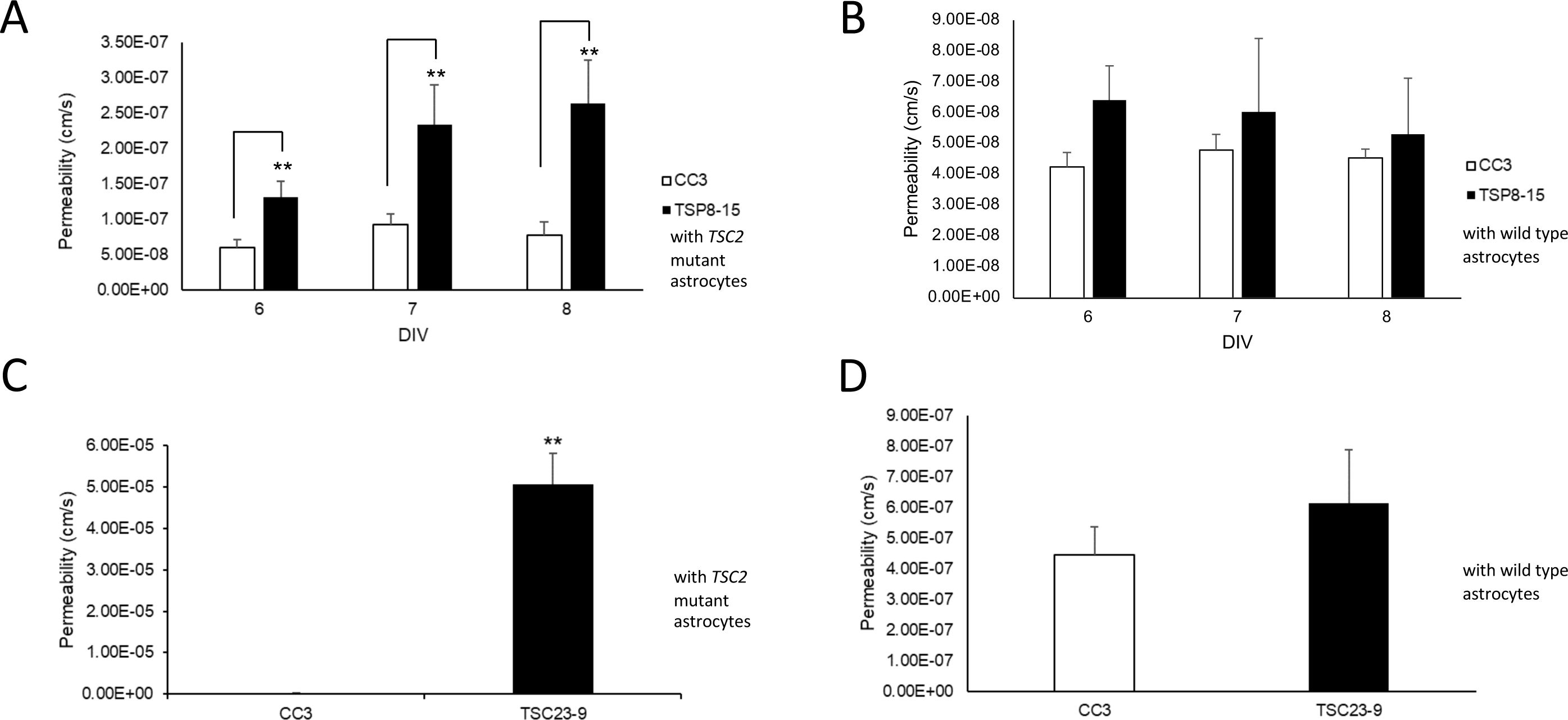
Control astrocytes rescue TSC BBB function. A) The BBB permeability measured on days 6-8 *in vitro* (DIV) in *TSC2* mutant (TSP8-15) NVUs is significantly higher than the one in control (CC3) NVUs (p < 0.01, N=5). **B)** The BBB permeability in *TSC2* (TSP8-15) mutant NVUs seeded with control (CC3) astrocytes is not significantly different from the BBB permeability measured in control (CC3) NVUs (N=5, p>0.05). **C)** The BBB permeability in NVUs seeded with BMECs, neurons and astrocytes derived from hiPSC derived from a different TSC patient (TSP23-9) carrying a distinct *TSC2* loss of function mutation is also significantly higher compared to control NVUs (CC3 = 2.95 x 10^-7^ cm/s). (p < 0.001, N=5). **D)** The presence of control (CC3) astrocytes in otherwise *TSC2* mutant (TSP23-9) NVUs rescued BBB permeabilities to levels statistically indistinguishable from that measured in control (CC3) NVUs. (p= N.S., N=5).

**Figure 7.**
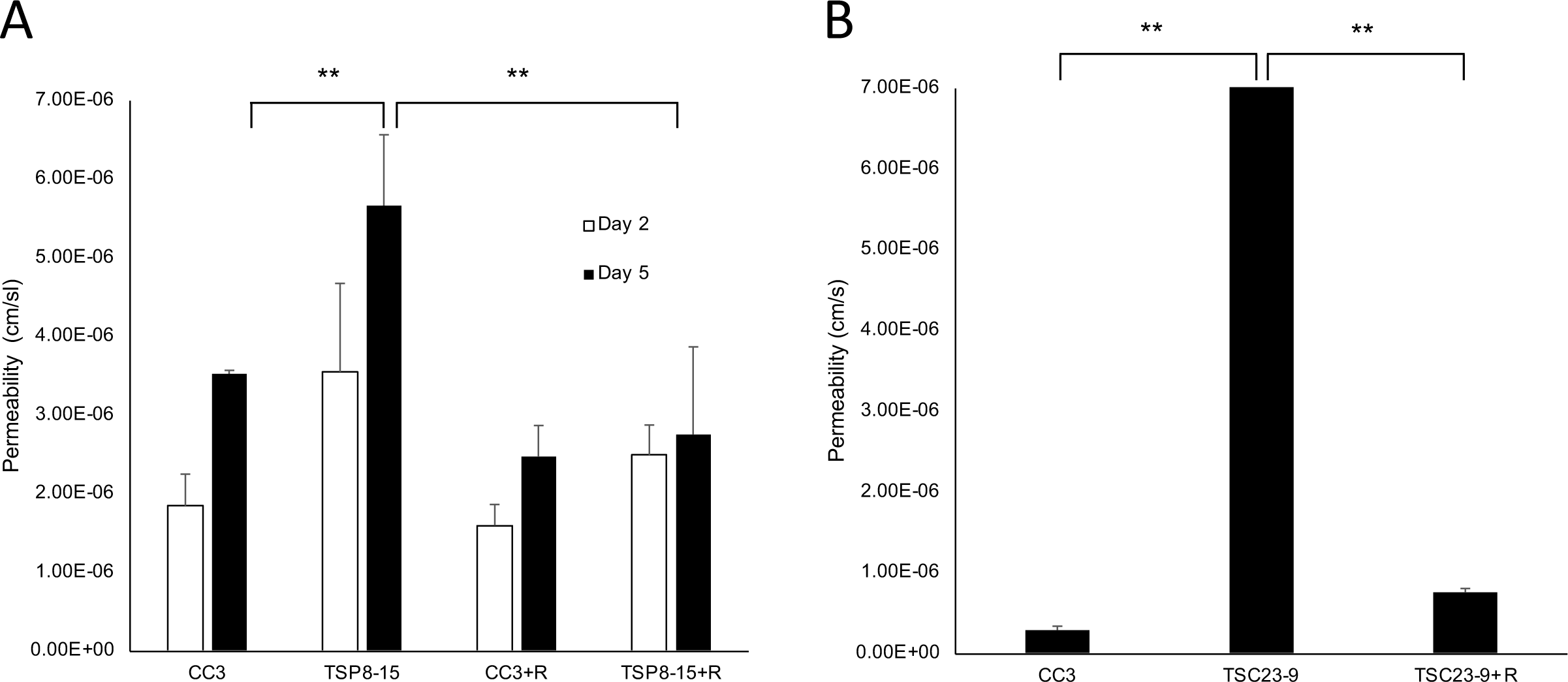
Rapamycin rescues TSC BBB permeability. A) On day 5 *in vitro*, BBB permeability of TSP8-15 NVUs is significantly higher than that seen in CC3 NVUs (p< 0.01; N=5). Perfusion of the vascular compartment with rapamycin results in TSC BBB permeabilities statistically significantly reduced compared to vehicle-treated TSC NVUs and statistically indistinguishable from controls (CC3). **B)** As observed for the TSP8-15 NVUs, the BBB permeability of TSP23-9 NVUs is significantly higher than in CC3 NVUs on 5 day in culture (p < 0.001, N=5). BBB permeabilities were significantly smaller in rapamycin perfused TSP23-9 NVUs than their vehicle treated counterparts and at levels statistically indistinguishable from CC3 levels. The error bar for the TSP23-9 sample is too small to be resolved on this graph (p N.S., N=5).

Given the heightened difference in BBB permeability in NVUs (containing BMECs, neurons and astrocytes) compared to the transwell cultures (only containing BMECs), we hypothesized that a cell type other than BMECs, such as astrocytes contribute to BBB function either directly or through regulation of BMECs. A crucial strength of our experimental approach is the ability to independently generate cell lineages (BMECs, astrocytes, neurons) of different genotypes and then load them in desired combinations (T*SC2*-wild type or – heterozygous mutant) into individual NVUs. To test the role that astrocytes play in the TSC BBB phenotype, we compared BBB permeabilities in NVUs comprised of *TSC2* mutant cells only and NVUs comprised of *TSC2* mutant cells except for the astrocytes which were *TSC2* wild type. This incorporation of control astrocytes into otherwise *TSC2* mutant NVUs rescued BMEC barrier function to levels similar to those measured in control NVUs at all time points measured (**Figure 6B**). The same observation was made in NVUs populated with BMECs, astrocytes, and neurons differentiated from another human iPSC line (TSP23-9) derived from a different patient with the *TSC2* gene harboring a distinct heterozygous *TSC2* mutation. As observed for TSP8-15, BBB permeability in the TSP23-9 NVU was increased compared to control NVU (**Figure 6C**). Importantly, permeability was also rescued by substituting *TSC2* wild type (CC3) astrocytes into otherwise TSP23-9 mutant NVU (**Figure 6D**). Collectively, these data indicate that control astrocytes are sufficient to rescue *TSC2* mutant impaired BBB permeability.

A central impact of *TSC2* gene mutations is dysregulation of mTOR kinase signaling. Application of rapamycin, an mTOR inhibitor, to the vascular compartment of the NVUs resulted in a significant decrease of the BMEC barrier permeabilities in TSC NVUs when compared to vehicle treated TSC NVUs, but did not affect BBB permeabilities in control NVUs (**Figure 7A, 7B**). We conclude that the impaired *TSC2*-mutant BBB function is reversible by a 24 hour treatment with rapamycin (10nM).

Given the impact of rapamycin on the TSC BBB phenotype, and the role mTOR signaling plays in the regulation of metabolism (Blenis 2017, Saxton and Sabatini 2017) we decided to investigate effects of genotype and rapamycin treatment on the exometabolome of the vascular and neural cell compartments of the NVU. We collected effluent from both the vascular and the neural cell side from control (CC3) and *TSC2*-mutant (TSP8-15) NVUs and used UPLC-IM-MS to perform unbiased metabolomics analysis. Not surprisingly we observed marked exometabolome differences between compartment types (neural cell versus vascular). Rapamycin treatment appeared to also cause clear differences in the metabolome, while the genotype (wild type versus heterozygous *TSC2* mutation) had a smaller effect as shown in a principal component analysis (PCA, **Figure 8A**) and hierarchical metabolite heatmap (**Figure 8B)**.

**Figure 8.**
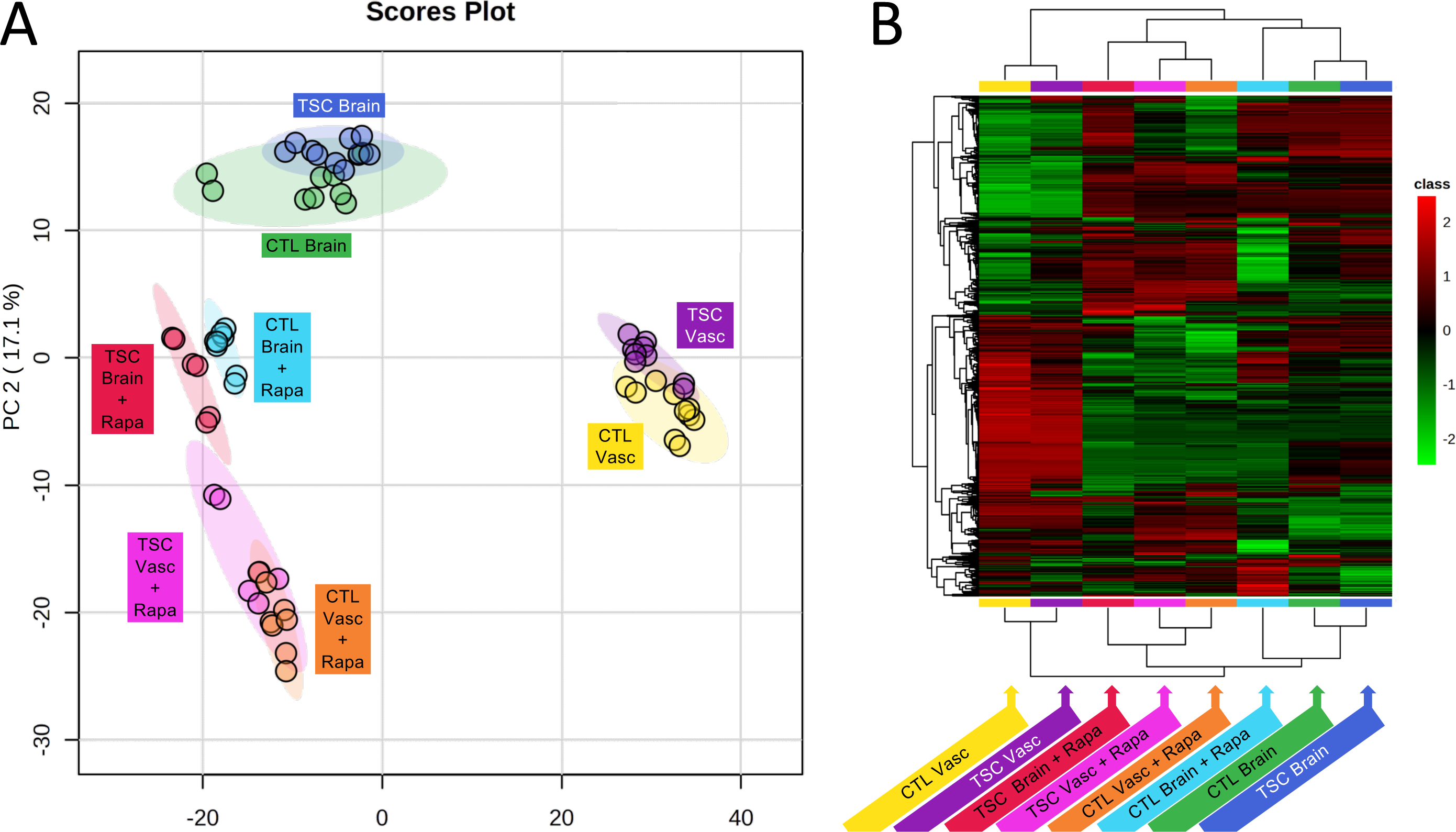
**Effect of genotype and rapamycin on neural cell and BBB NVU exometabolomes**. **A)** Principal Component Analysis indicates a pronounced metabolic difference between tissue types (vascular versus brain). In addition, rapamycin treatment for brain and vascular compartments causes a shift in the metabolome. The genotype (wild type, CC3 versus *TSC2* heterozygous mutation, TSP8-15) appeared to affect the metabolome to a much lesser degree. **B)** Hierarchical clustering maps provide global comparisons for differences of metabolite levels > 2-fold with a p value of less than 0.05. Different individual metabolite levels (rows) are clustered by genotype, tissue type and rapamycin treatment (columns). Metabolites are colored according to relative feature abundance across all samples ranging from low (green) to high (red).

We performed a pathways analysis and show the top 25 pathways that differed between genotype (wild type versus *TSC2* heterozygous mutant) and rapamycin treatment (**Figure 9**). We identified 5 pathways that differed between wild type and *TSC2*-mutant vascular tissue, these included caffeine-, purine-, sialic acid-, vitamin B6– and tyrosine metabolism. The same comparison between wild type and *TSC2*-mutated neural compartment revealed 3 different pathways (prostaglandin synthesis, linoleate– and pyrimidine metabolism). Rapamycin affected a total of 10 and 9 pathways in the vascular and neural compartments, respectively. Four pathways were affected in both tissues, they include urea cycle/amino group-, glutamate-, tryptophan– and histidine metabolism (**Figure 9**).

**Figure 9.**
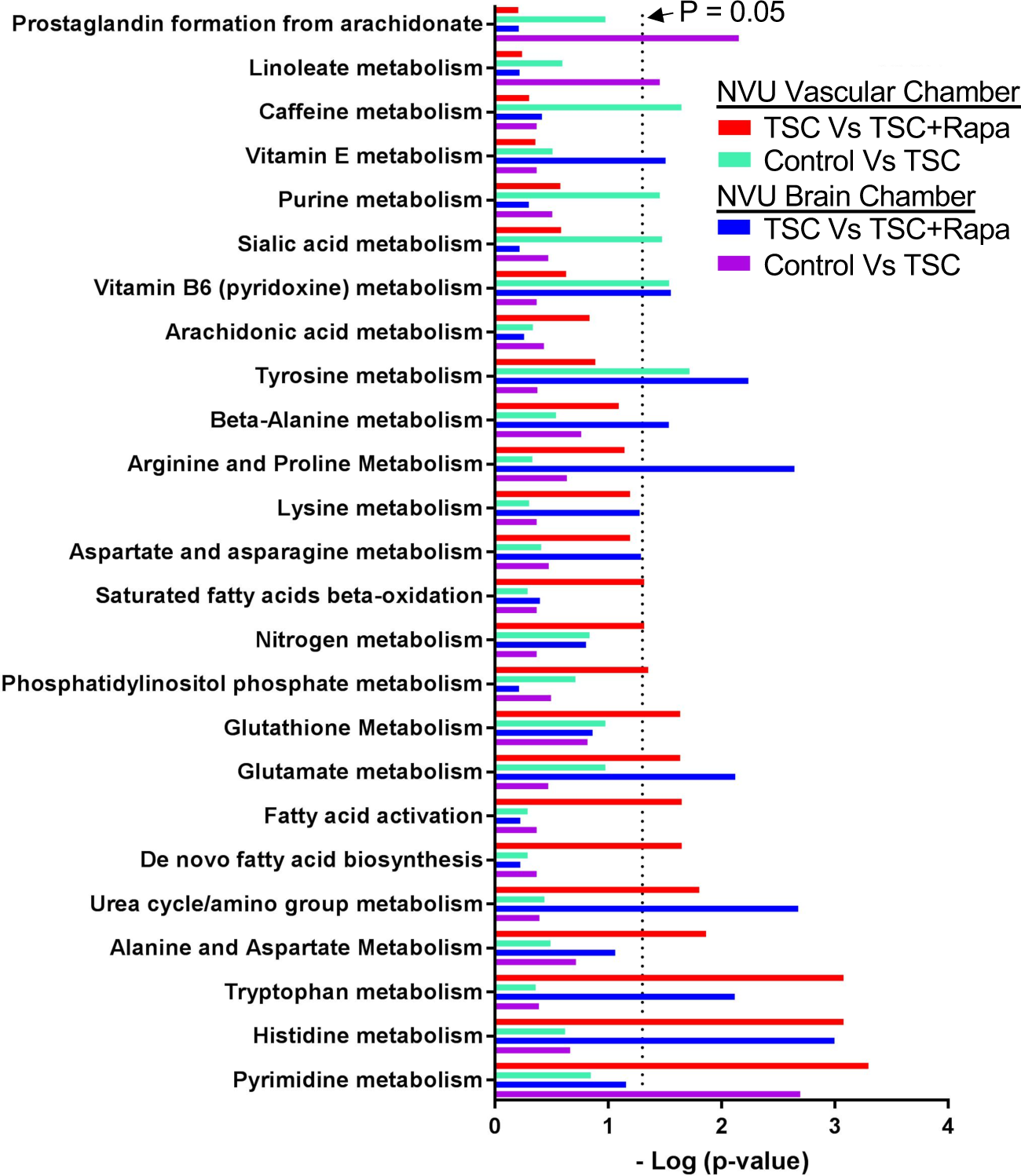
Metabolic pathways altered by rapamycin, genotype, and tissue type in control (CC3) and TSC (TSP8-15) NVUs. The top 25 significant response pathways are shown. In each pathway listed, at least one variable (rapamycin treatment, genotype, tissue type) showed a significant difference. Significance was defined as p-values less than or equal to 0.05 in pathways that had four or more metabolites with altered expression levels at a minimal two-fold change.

Finally, we used RNA Seq to look at differences in gene expression between control (CC3) *TSC2* mutant (TSP8-15) neural cell compartments. In a comparison of TSP8-15 versus control (CC3), a total of 1143 up-regulated and 769 down-regulated genes were identified (log2foldchange >1 and padj<0.05). The top 100 most differentially expressed genes are listed in **Supplementary** Figure 4. GO pathways analysis identified multiple pathways pertaining to synapse function and formation, and glutamatergic synapses in particular. In addition, oxidative phosphorylation, ATP synthesis coupled electron chain transport, and respiratory electron chain transport, all pertaining to cellular respiration are also different in wild type and *TSC2*-heterozygus mutant neural compartments (**Table 1**).

**Table 1.**
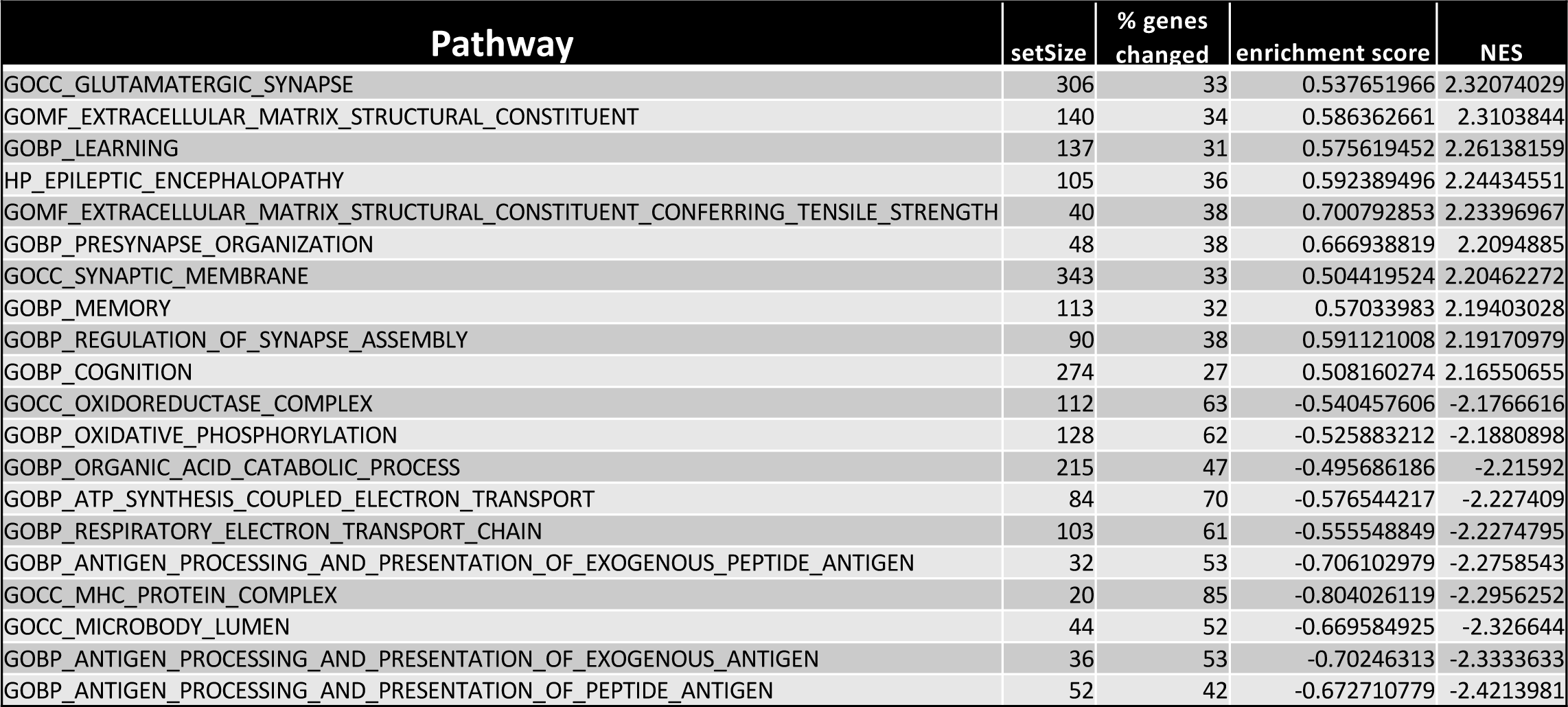
Top 10 up– and down regulated signaling pathways in TSC neural compartment when compared to control. Based on the normalized enrichment score (NES) we analyzed RNA Seq data and rank the top 10 upregulated and the top 10 down regulated pathways in a comparison of *TSC2* mutant versus control neural cell chambers.

| Pathway | setSize | % genes changed | enrichment score | NES |
| --- | --- | --- | --- | --- |
| GOCC_GLUTAMATERGIC_SYNAPSE | 306 | 33 | 0.537651966 | 2.32074029 |
| GOMF_EXTRACELLULAR_MATRIX_STRUCTURAL_CONSTITUENT | 140 | 34 | 0.586362661 | 2.3103844 |
| GOBP_LEARNING | 137 | 31 | 0.575619452 | 2.26138159 |
| HP_EPILEPTIC_ENCEPHALOPATHY | 105 | 36 | 0.592389496 | 2.24434551 |
| GOMF_EXTRACELLULAR_MATRIX_STRUCTURAL_CONSTITUENT_CONFERRING_TENSILE_STRENGTH | 40 | 38 | 0.700792853 | 2.23396967 |
| GOBP_PRESYNAPSE_ORGANIZATION | 48 | 38 | 0.666938819 | 2.2094885 |
| GOCC_SYNAPTIC_MEMBRANE | 343 | 33 | 0.504419524 | 2.20462272 |
| GOBP_MEMORY | 113 | 32 | 0.57033983 | 2.19403028 |
| GOBP_REGULATION_OF_SYNAPSE_ASSEMBLY | 90 | 38 | 0.591121008 | 2.19170979 |
| GOBP_COGNITION | 274 | 27 | 0.508160274 | 2.16550655 |
| GOCC_OXIDOREDUCTASE_COMPLEX | 112 | 63 | -0.540457606 | -2.1766616 |
| GOBP_OXIDATIVE_PHOSPHORYLATION | 128 | 62 | -0.525883212 | -2.1880898 |
| GOBP_ORGANIC_ACID_CATABOLIC_PROCESS | 215 | 47 | -0.495686186 | -2.21592 |
| GOBP_ATP_SYNTHESIS_COUPLED_ELECTRON_TRANSPORT | 84 | 70 | -0.576544217 | -2.227409 |
| GOBP_RESPIRATORY_ELECTRON_TRANSPORT_CHAIN | 103 | 61 | -0.555548849 | -2.2274795 |
| GOBP_ANTIGEN_PROCESSING_AND_PRESENTATION_OF_EXOGENOUS_PEPTIDE_ANTIGEN | 32 | 53 | -0.706102979 | -2.2758543 |
| GOCC_MHC_PROTEIN_COMPLEX | 20 | 85 | -0.804026119 | -2.2956252 |
| GOCC_MICROBODY_LUMEN | 44 | 52 | -0.669584925 | -2.326644 |
| GOBP_ANTIGEN_PROCESSING_AND_PRESENTATION_OF_EXOGENOUS_ANTIGEN | 36 | 53 | -0.70246313 | -2.3333633 |
| GOBP_ANTIGEN_PROCESSING_AND_PRESENTATION_OF_PEPTIDE_ANTIGEN | 52 | 42 | -0.672710779 | -2.4213981 |

Gene set analysis revealed an upregulation of K-ras signaling (**Supplementary** Figure 5A), glutamatergic synapse (**Supplementary** Figure 5B), and epileptic encephalopathy (**Supplementary** Figure 5C) when comparing *TSC2* mutant (TSP8-15) versus control (CC3) neural cells for RNA expression differences.

## Discussion

Neurogenetic disorders underlie many if not most causes of epilepsy, autism, and intellectual disabilities in children. TSC is a prototypical disease for understanding these severe manifestations due to known genetic causes, relevant downstream signaling pathways, and FDA-approved medications to treat some aspects of the disorder. TSC-associated epilepsy is one of the most debilitating aspects given the highly prevalent nature of seizures that are often very hard to control using standard approaches. Many patients in fact require multiple anti-seizure medications and consideration of dietary and surgical therapies.

Assessing mechanism underlying these pathologies has been done in animal model of TSC, and we have extensively modeled TSC using zebrafish and mice (Uhlmann, Wong et al. 2002, Wong, Ess et al. 2003, Kim, Speirs et al. 2011, Carson, Van Nielen et al. 2012, Carson, Fu et al. 2013, Carson, Kelm et al. 2015). However, assessing cell-type specific contributions requires modifying the genotype of specific neural cell lineages which is very challenging. Here we use combinatorial human cell-based systems which present a unique opportunity to identify the contribution of multiple CNS cell lineages to the formation of the epileptic neural networks in TSC.

Genotype/phenotype relationships and gene dosage are key aspects in the pathogenesis of TSC. We chose to use *TSC2* mutant iPSCs as the basis for all patient-derived cells in this study. This reflects the much more frequent occurrence of *TSC2* versus *TSC1* mutations in patients with TSC, as well as the generally more severe manifestations seen in patients with *TSC2* mutations (Au, Williams et al. 2007). Importantly, all mutant cells we used here and showed a defect in the BBB were *TSC2* heterozygous. This begins to address a central question in the field: is a homozygous mutation required for disease manifestations? *TSC2* homozygous mutations were originally thought to be due to loss of heterozygosity (LOH) with “second hits” being acquired somatically during brain development and post-natal life. This model came from the initial assumption that the *TSC1* and *TSC2* gene would follow classic tumor suppressor paradigms including LOH. This concept was further supported by the many rodent and zebrafish models we and others have made over the past 10 years (Zeng, Rensing et al. 2011, Carson, Van Nielen et al. 2012, Fu and Ess 2013, Kim, Kowalski et al. 2013, Carson, Kelm et al. 2015). In these animal models, heterozygous loss of *Tsc1* or *Tsc2* genes generally have very mild or no phenotypes, whereas homozygous *Tsc1* or *Tsc2* mutant animals usually have severe manifestations. This contradiction between patient-based data from deep DNA sequencing of tubers as compared to animal models was a strong rationale for use of human iPSCs here. The minimal evidence for second hit homozygous mutant cells in the human brain has led to suggestions that heterozygous mutant cells are sufficient to cause disease in humans. More recent work in human brain and organoids (Giannikou, Lasseter et al. 2021, Eichmuller, Corsini et al. 2022) further suggests that LOH may not be required, especially in cells that natively have lower levels of hamartin or tuberin protein. Other groups have presented data that supports a role for heterozygous as well as homozygous *TSC2* mutations in the pathogenesis of TSC (Blair, Hockemeyer and Bateup 2018, Winden, Sundberg et al. 2019).

Our studies show that astrocytes play a crucial role in the observed TSC BBB phenotype since control wild type astrocytes were able to rescue the TSC BBB dysfunction. We also recently reported impaired glutamate uptake by TSC as compared to wild type human astrocytes derived from the same iPSC cell lines used here (Miller, Schaffer et al. 2021). Thus, the findings reported here further highlight the critical role of astrocytes in TSC pathogenesis. Prior work in mouse models also implicated astrocyte dysfunction but mainly focused on homozygous mutant cells (Wong, Ess et al. 2003, Zeng, Ouyang et al. 2007). These findings are important, as they emphasize species-specific differences of genotype upon rodent versus human cell-based models of TSC and other related disorders. Another recent report details abnormalities of the mouse BBB and connection to seizures in an astrocyte restricted homozygous knockout of the *Tsc1* gene (Guo, Zhang et al. 2023). Interestingly, treatment with RepSox, a TGF-β inhibitor that stabilizes the BBB decreased seizures and increased survival in this well-established mouse model of TSC.

Using metabolomics and RNA Seq, our results further detail changes in many pathways that are impacted by genotype and cell type. Notably, *TSC2* heterozygous mutant BBB versus control BBB did not show increased mTOR signaling (data not shown). This is not unexpected, as human cells with heterozygous mutations generally do not show increased mTOR signaling using conventional immunoblotting, though homozygous mutant human cells usually do. Small differences in mTORC1 signaling from human *TSC2* heterozygous cells with the very low amount of protein we can extract from an individual NVU however may not be detectable with the sensitivity of current methods. We did see an impact of genotype and rapamycin treatment on the metabolome of both the vascular and brain compartments of the NVU (**Figures 8 and 9**). Modest alterations in mTOR signaling then seem to be present given the impact of low dose rapamycin. This suggests that patients with TSC have diffusely abnormal BBB as all cells in patients are thought to be at least heterozygous mutant. This may be supported by clinical imaging studies in patients with TSC showing altered Apparent Diffusion Coefficient (ADC) brain imaging in TSC (Garaci, Floris et al. 2004). Other studies show increased inflammatory markers in the brains of patients with TSC, further supporting a potential mechanism of a “leaky” BBB allowing ingress of immune cells to the brain furthering dysfunction (Boer, Jansen et al. 2008, Boer, Troost et al. 2008). Alternatively, the abundance of these inflammatory markers may result from seizures secondarily causing BBB dysfunction. This further validates our approach with human iPSC and microphysiological systems as cell/cell functional interactions can be defined without confounding from seizures seen in most TSC animal models.

Our PCA analysis revealed greater metabolome differences between neural cell and vascular compartments and rapamycin treatment versus vehicle than between *TSC2* genotypes. While there is chemical communication across the BBB, the major contributor to metabolic differences between the vascular and neural compartments are likely metabolic differences in cell lineages (BMECs versus neuron and astrocytes) involving many additional cellular processes aside from signaling pathways associated with *TSC2*. We identified a greater number of metabolic pathways changed by rapamycin than affected by the *TSC2* genotype in the vascular– and neural compartments. Assessment of RNA transcriptional profiles of *TSC2* wild type and TSC2-mutated neural compartments revealed differentially regulated pathways associated with synapse development and mitochondrial respiration, cellular functions intricately linked to CNS development.

## Conclusions

In summary, we found altered BBB function within a human *TSC2* heterozygous NVU microphysiological system. Replacement of *TSC2*-mutated astrocytes with *TSC2* wild type astrocytes or treatment with rapamycin was sufficient to rescue the BBB phenotype. These findings have translational and clinical impact as impaired BBB may contribute to neurological disorders and future therapeutics for TSC may be designed to improve BBB function. As rapamycin and related compounds are being used clinically for some aspects of TSC, it is possible that alteration of the BBB permeability may be occurring and further focus on the BBB should yield important data and insights that may have clinical impact.

## Declarations

- Ethics approval and consent to participate-all human samples were obtained under IRB approved studies at Vanderbilt University Medical Center (#080369) or Boston Children’s Hospital (#P00008224).
- Consent for publication-Not applicable
- Availability of data and materials-available upon request
- Competing interests-Dr. Mustafa Sahin reports grant support from Novartis, Biogen, Astellas, Aeovian, Bridgebio, and Aucta. He has served on Scientific Advisory Boards for Novartis, Roche, Regenxbio, SpringWorks Therapeutics, Jaguar Therapeutics, and Alkermes.
- Funding-Research reported in this publication was supported in part by the National Center for Advancing Translational Sciences of the National Institutes of Health under Award No. UH3TR002097 and, through Vanderbilt University Medical Center, Award No. UL1TR002243 (K.C.E., J.P.W., M.D.N.). Also supported by the National Institutes of Health (R01NS118580 (R.A.I. and K.C.E.), and U54NS092090 (M.S.). The content is solely the responsibility of the authors and does not necessarily represent the official views of the National Institutes of Health.
- Authors’ contributions-JAB planned and performed experiments, analyzed data, wrote the manuscript; SLF performed experiments, analyzed data, wrote the manuscript; MJ, performed experiments, analyzed data, wrote the manuscript; PW performed experiments, analyzed data, wrote the manuscript; RAI planned experiments, analyzed data, wrote the manuscript, provided funding; RC analyzed data, wrote the manuscript; LA performed experiments, wrote the manuscript; MS, analyzed data, wrote the manuscript; JPW planned experiments, analyzed data, wrote the manuscript, provided funding; KCE planned experiments, analyzed data, wrote the manuscript, provided funding; MDN planned and performed experiments, analyzed data, wrote the manuscript, provided funding.

## Abbreviations

ANOVA: ANalysis Of Variance
BBB: Blood-Brain Barrier
BMEC: Brain Microvascular Endothelial-like Cells
iPSC: induced Pluripotent Stem Cell
GSEA: Gene Set Enrichment Analysis
LCHRMS: Liquid Chromatography-High Resolution Mass Spectrometry
mTOR: mechanistic/mammalian Target Of Rapamycin
NVU: NeuroVascular Unit
PCA: Principal Component Analysis
TEER: TransEndothelial Electrical Resistance
TSC: Tuberous Sclerosis Complex

## Acknowledgements

This work was supported in part using the resources of the Center for Innovative Technology at Vanderbilt University. The microfluidic NVUs were fabricated by David K. Schaffer and Clayton M. Britt of the Vanderbilt Microfabricated Technologies Resource. The data that support the findings of this study are available from the corresponding author upon reasonable request. The Vanderbilt Creative Data Solutions Shared Resource (RRID:SCR_022366) performed the RNA-Seq data processing, differential gene expression analysis and Gene Set Enrichment Analysis.

## Figure Legends

**Supplementary Figure 1. iPSC Karyotypes.** Karyotypes of all cell lines used were normal.

**Supplementary Figure 2. Cortical neuronal cultures differentiated from control– and TSC-iPSC lines express the neuronal markers** β**3-tubulin and MAP2.** Cortical neuronal cultures differentiated from control iPSC lines (**A**) and TSC iPSC lines (**B**) show a dense network of β3-tubulin– and MAP2-positive neurites. The fluorescence signals of the images in Fig. 2 were enhanced for optimal visualization of the markers; therefore, comparisons of expression levels between the different iPSC lines are not possible from this Figure. We have not observed any obvious and consistent genotype-dependent difference in the level or distribution of β3-tubulin or MAP2 expression. TSP77+/+ and TSP77+/− are isogenic TSC wild type and TSC2 heterozygous mutated iPSC lines, respectively. The neuronal cultures shown here were differentiated between 67-104 days before seeding into the NVUs. Scale bar is 100 μm.

**Supplementary Figure 3. Astrocyte cultures differentiated from control and TSC-iPSC lines express the astrocytic markers GFAP and S100B.** Astrocyte cultures differentiated from control iPSC lines and TSC iPSC lines show expression of GFAP and S100B. The expression levels of both GFAP and S100B varies from cell to cell within the same culture. A few cells express detectable levels of GFAP, but no S100B, some cells express S100B, but no GFAP and the majority of cells express both markers. Generally, the expression of S100B in control and TSC astrocytes co-expressing GFAP is lower than in cells expressing S100B only. The signals of the images were enhanced for optimal visualization of the markers and cell morphology; therefore, comparisons of expression levels between the lines are not possible from this Figure. However, we have not observed any obvious and consistent differences in GFAP or S100B expression between genotypes. TSP77+/+ and TSP77+/− are isogenic TSC2 wild type and TSC2 heterozygous mutated iPSC lines, respectively. The astrocyte cultures shown here were differentiated for >200 days. Scale bar is 100 μm.

**Supplementary Figure 4. Genes expressed differentially in control (CC3) and TSPC-patient (TSP8-15) derived NVU neural chambers.** The top 100 genes sorted by fold-difference with increased (blue-green) or decreased (orange) expression levels in the neural chamber of TSC2 mutant (TSP-15) compared to control (CC3) are shown.

**Supplementary Figure 5. Gene set analysis of three gene pathways potentially relevant to TSC pathogenesis.** We provide gene set analysis for three pathways that might be relevant to TSC CNS pathogenesis: K Ras signaling (**A,** NES = 1.69, setSize 178), Glutamatergic Synapse (**B**, NES = 2.32, setSize: 306) and Epileptic Encephalopathy (**C**, NES = 2.26, setSize 105).

