## Supplemental Figures for "Rescue of Impaired Blood-Brain Barrier in Tuberous Sclerosis Complex Patient Derived Neurovascular Unit"

**Suppl. Figure 1: iPSC line Karyotypes**

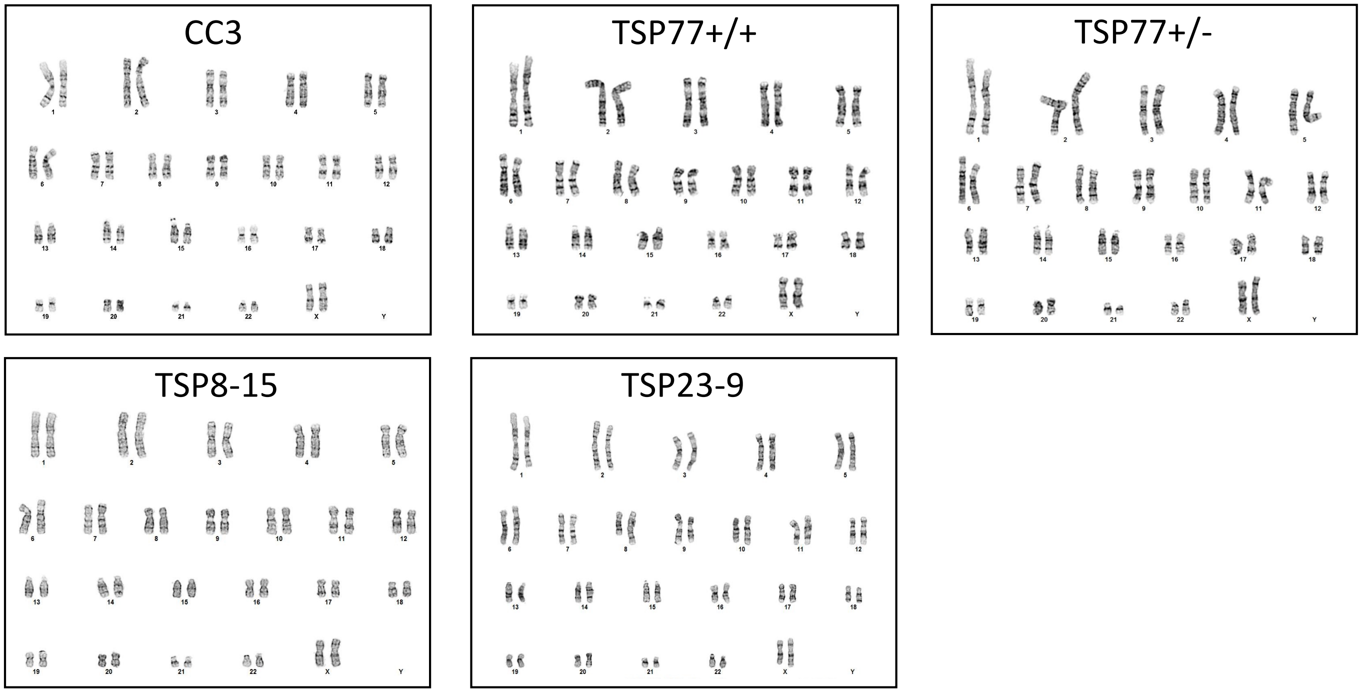

### Suppl. Figure 2A: Neuronal marker expression following differentiation of control iPSC lines

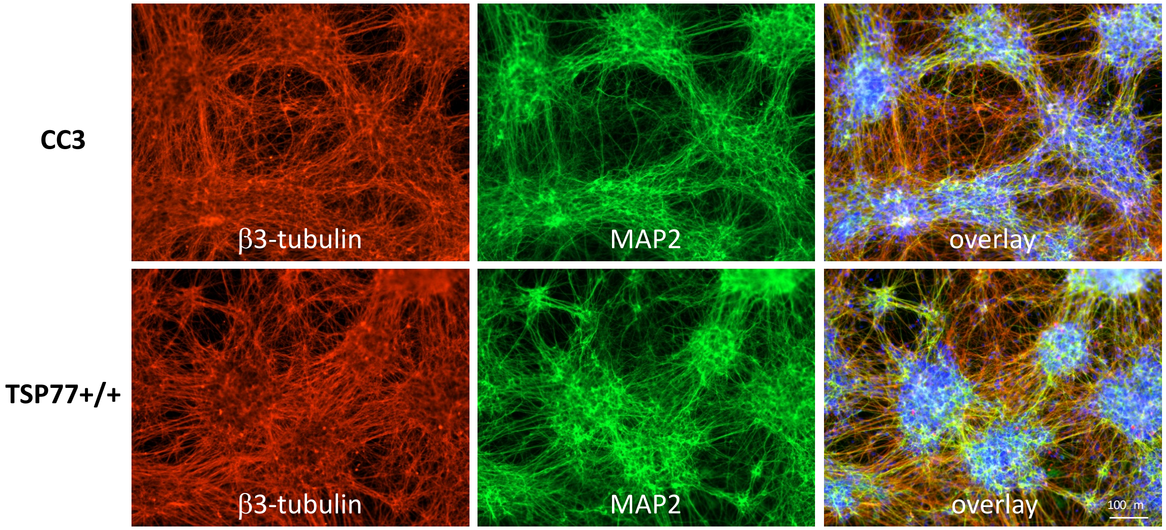


### Suppl. Figure 2 Legend:

**Suppl. Fig. 2. Cortical neuronal cultures differentiated from control- and TSC-hiPSC lines express the neuronal markers β3-tubulin and MAP2.** Cortical neuronal cultures differentiated from control hiPSC lines (**A**) and TSC hiPSC lines (**B**) show a dense network of β□-tubulin- and MAP2-positive neurites. The fluorescence signals of the images in Fig. 2 were enhanced for optimal visualization of the markers; therefore comparisons of expression levels between the different hiPSC lines are not possible from this Figure. We have not observed any obvious and consistent genotype-dependent difference in β3-tubulin or MAP2 expression. TSP77+/+ and TSP77+/- are isogenic TSC wildtype and TSC2 heterozygous mutated hiPSC lines, respectively. The cultures shown here were differentiated between 67-104 days. Scale bar is 100 □m.

**Suppl. Figure 2B: Neuronal marker expression following differentiation of TSC iPSC lines**

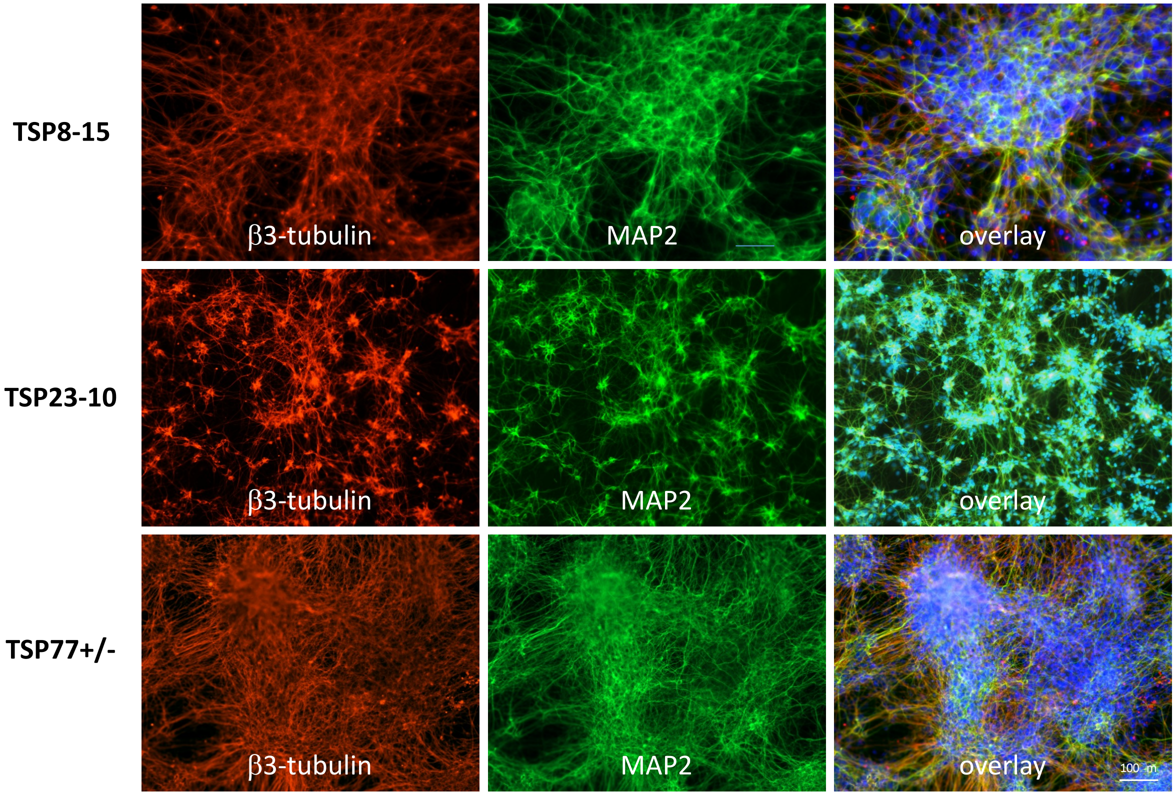

**Suppl. Figure 3: Expression of astrocyte markers by astrocytes differentiated from control and TSC hiPSC lines**

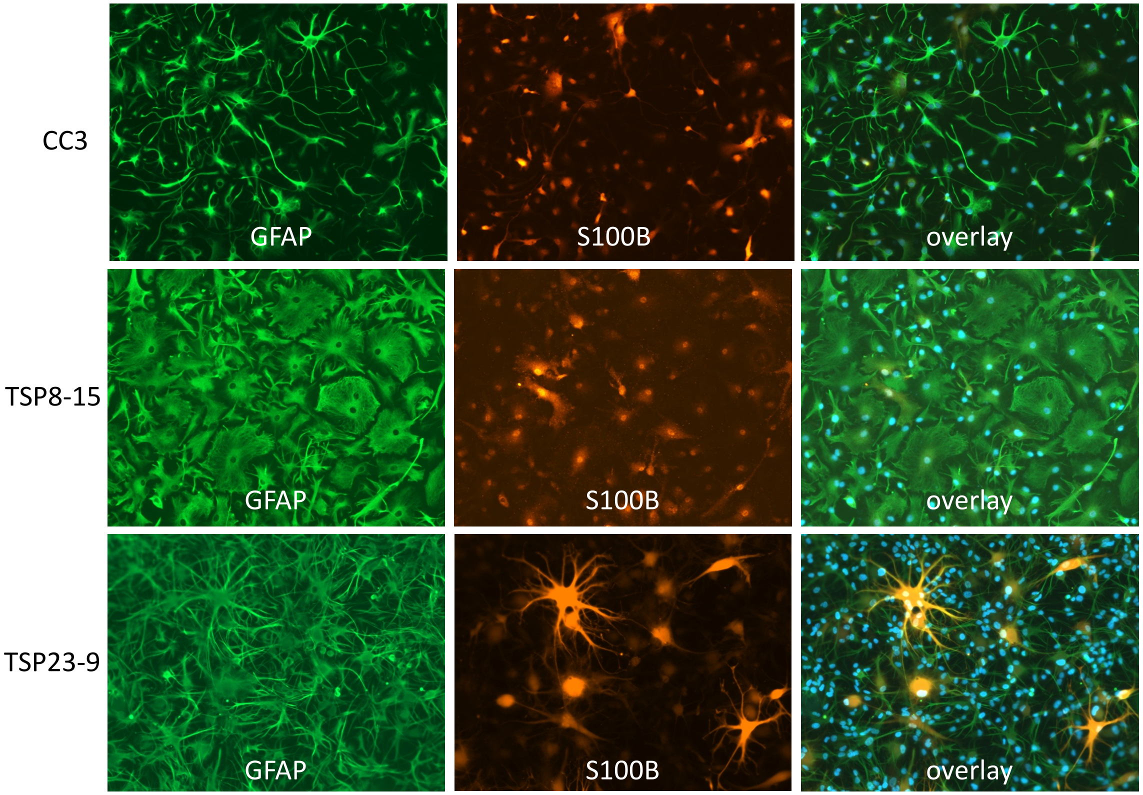

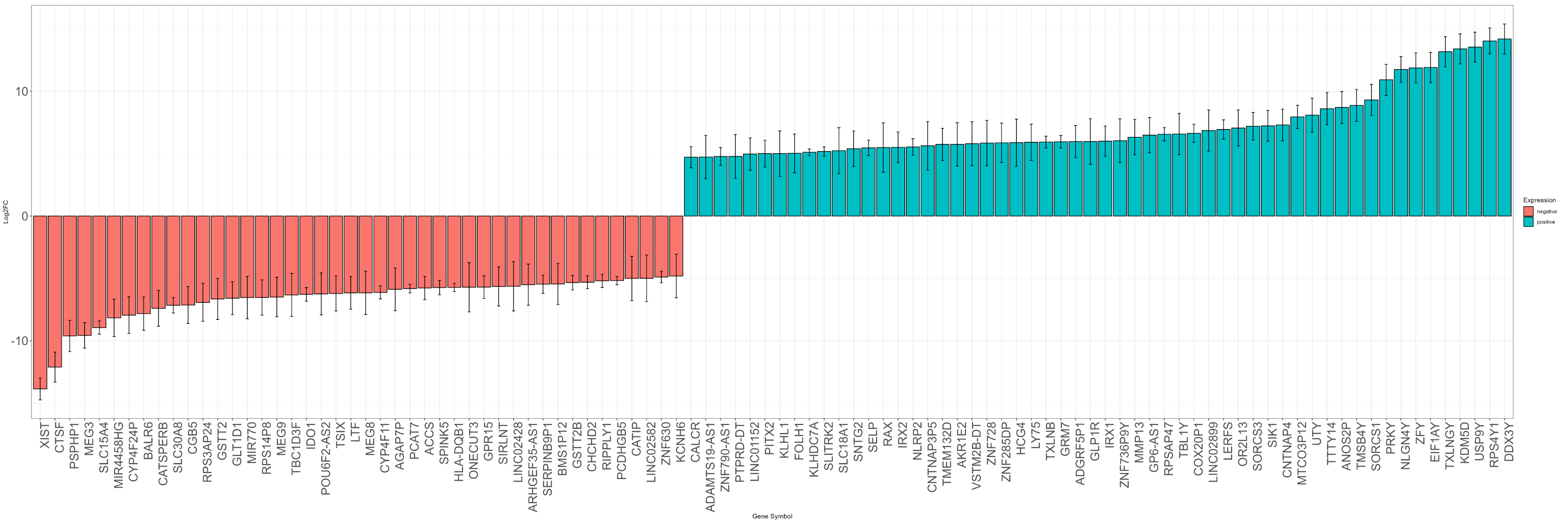


**Supplementary Figure 4.**

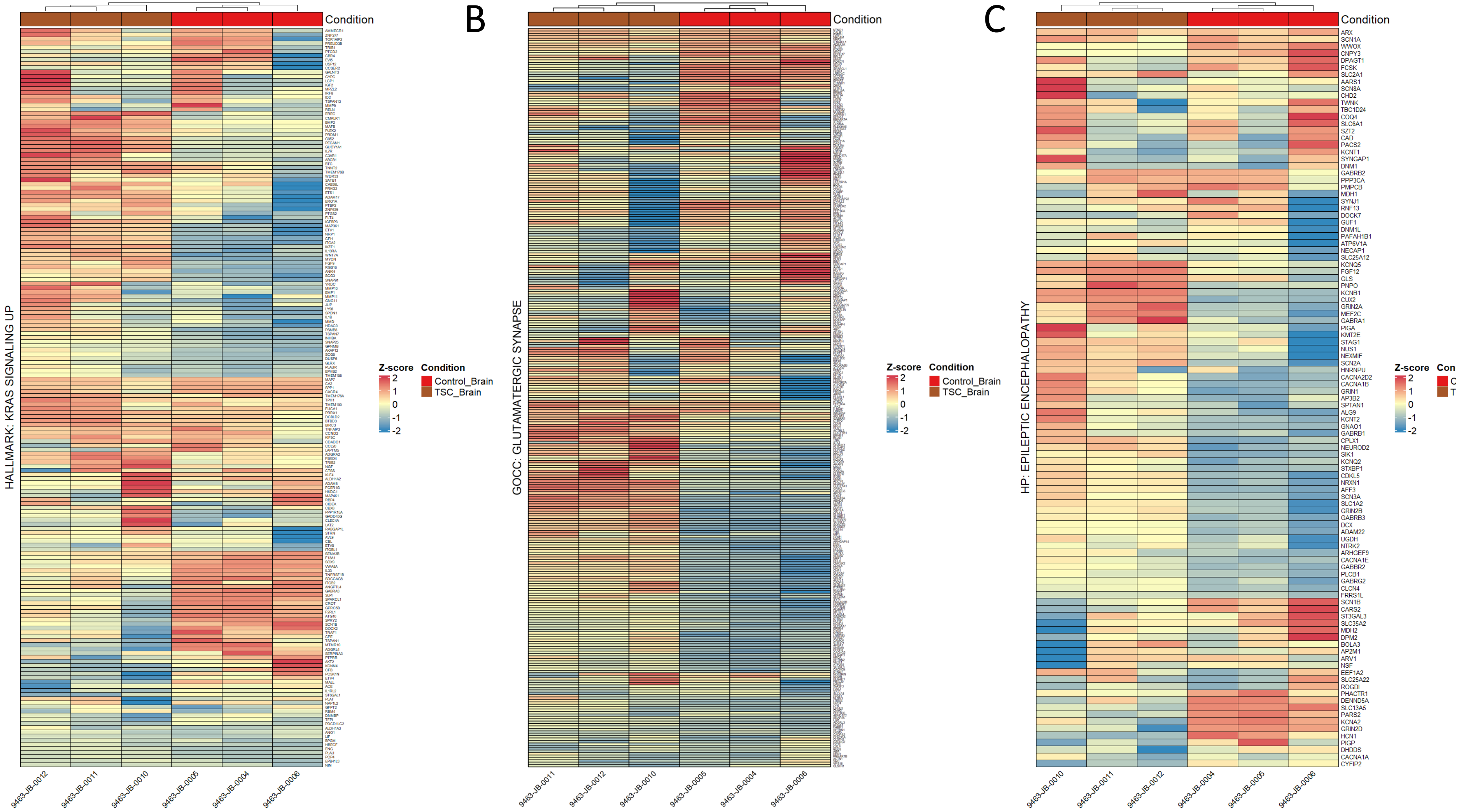
B
C
Condition
HALLMARK: KRAS SIGNALING UP
GOCC: GLUTAMATERGIC SYNAPSE
HP: EPILEPTIC ENCEPHALOPATHY
Z-score
2
1
0
-1
-2
Condition
Control_Brain
TSC_Brain
9463-JB-0012
9463-JB-0011
9463-JB-0010
9463-JB-0005
9463-JB-0004
9463-JB-0006
ARX
SCN1A
WWOX
CNPY3
DPAGT1
FCSK
SLC2A1
AARS1
SCN8A
CHD2
TWNK
TBC1D24
COQ4
SLC6A1
SZT2
CAD
PACS2
KCNT1
SYNGAP1
DNM1
GABRB2
PPP3CA
PMPCB
MDH1
SYNJ1
RNF13
DOCK7
GUF1
DNM1L
PAFAH1B1
ATP6V1A
NECAP1
SLC25A12
KCNQ5
FGF12
GLS
PNPO
KCNB1
CUX2
GRIN2A
MEF2C
GABRA1
PIGA
KMT2E
STAG1
NUS1
NEXMIF
SCN2A
HNRNPU
CACNA2D2
CACNA1B
GRIN1
AP3B2
SPTAN1
ALG9
KCNT2
GNAO1
GABRB1
CPLX1
NEUROD2
SIK1
KCNQ2
STXBP1
CDKL5
NRXN1
AFF3
SCN3A
SLC1A2
GRIN2B
GABRB3
DCX
ADAM22
UGDH
NTRK2
ARHGEF9
CACNA1E
GABBR2
PLCB1
GABRG2
CLCN4
FRRS1L
SCN1B
CARS2
ST3GAL3
SLC35A2
MDH2
DPM2
BOLA3
AP2M1
ARV1
NSF
EEF1A2
SLC25A22
ROGDI
PHACTR1
DENND5A
SLC13A5
PARS2
KCNA2
GRIN2D
HCN1
PIGP
DHDDS
CACNA1A
CYFIP2
